# Prey abundance drives habitat occupancy by jaguars in Amazonian floodplain river islands

**DOI:** 10.1101/555052

**Authors:** Rafael M. Rabelo, Susan Aragón, Júlio César Bicca-Marques

## Abstract

The jaguar (*Panthera onca*) is widely distributed across a broad range of habitat types, where its feeding habits and habitat use patterns vary significantly. The jaguar and its main arboreal prey – the brown-throated sloth (*Bradypus variegatus*) and the red howler monkey (*Alouatta juara*) – are widespread in the Amazonian floodplain forests of the Mamirauá Reserve. These forest-dwelling species are the most common mammal species both in the continuous forest and the forest patches surrounded by a river matrix – the fluvial islands – of the Solimões and Japurá rivers. We used sign surveys along line-transects to assess the pattern of habitat occupancy by jaguars in Amazonian floodplain forests. Specifically, we (i) tested whether habitat occupancy by jaguars differs between river islands and continuous forest; and (ii) evaluated whether and how the local abundance of sloths and howler monkeys influence the probability of site occupancy by jaguars. We built an occupancy model and used Bayesian inference to reach these goals. The proportion of sites estimated to be used by jaguars was *ψ* = 0.75 (HPD95: 0.36–1.00), and it did not differ between islands and continuous forest. The abundance of both prey species had a direct influence on jaguar’s habitat use, whereas the aquatic matrix seems to have a negligible effect on the use of islands by jaguars. We conclude that the isolation of the river islands within the aquatic matrix does not hamper jaguars to use them. We also conclude that prey search modulates jaguars’ habitat occupancy patterns with both prey species having a similar effect. This finding is compatible with the previously reported importance of sloths to the diet of jaguars in the study region despite its lower abundance than howlers. Finally, we suggest that sign surveys are an alternative method to assess the pattern of jaguar habitat occupancy in floodplain forests.

## Introduction

Predation is a remarkable interspecific interaction that has long interested ecologists (Gause et al., 1936). Large carnivores are prominent top predators that prevent prey populations to overcrowd and deplete their food sources, and whose demise can initiate substantial cascading ecological effects in the food chain that compromise ecosystem structure and functioning (Ripple et al., 2014). Therefore, assessments of carnivore distribution and population size are essential for developing informed conservation actions for these species and their ecosystems.

Jaguars (*Panthera onca*) are the largest American felids. They are widely distributed across a broad range of habitat types (Sanderson et al., 2002), where their feeding habits and habitat use patterns vary significantly (Astete et al., 2007; Morato et al., 2016). They are opportunistic predators that exploit most of their medium to large terrestrial prey species (González and Miller, 2002) according to their availability (Rabinowitz and Nottingham, 1986). The species is often range resident and move over long daily distances (2.3–16.4 km) in highly variable home ranges (8.8–718.6 km^2^; Morato et al., 2016).

Jaguars’ predominant terrestriality does not preclude them from occurring in Amazonian seasonally flooded forests (herein *várzea* forests). This is the case at the Mamirauá Sustainable Development Reserve – a protected area of *várzea* forests in Central Amazon – where jaguars reach high densities probably because of high prey abundance (Ramalho, 2012). The availability of arboreal (e.g., sloths and monkeys; Rabelo et al., 2017) and water-associated (e.g., caimans and their eggs; Ramalho, 2012; Torralvo et al., 2017) prey species allows jaguars to reside yearlong in flooded forests, including the 4- to 6-month-long high-water season (Ramalho et al., 2009; Ramalho, 2012).

Red howler monkeys (*Alouatta juara*), brown-throated sloths (*Bradypus variegatus*) and jaguars are the most common mammal species in both the continuous forest and the fluvial islands (Rabelo et al., 2019). These islands originate from a complex river dynamics, in which the deposition of sediments as sandbars in the river channel are followed by primary succession (Kalliola et al., 1991). Therefore, island colonization by howlers, sloths and jaguars requires that they move through other land cover types, including the crossing of inhospitable water bodies by swimming (see Holt, 1932; Nunes, 2014).

Several factors influence the success of this dispersal through the matrix, including the distance to be crossed, the matrix permeability to the species movement, and the disperser’s motivation to find food resources in the new habitat (Lima and Zollner, 1996). Therefore, given that jaguars can use and easily move through a wide range of land cover types (Morato et al. 2018), it is plausible to expect that the search for prey plays a major role in their dispersal across rivers to reach fluvial islands.

Here we developed an occupancy modelling study of the pattern of habitat occupancy of Amazonian *várzea* forests by jaguars to test the influence of prey availability on it. We used jaguar sign surveys along line-transects together with records of two important prey species, the brown-throated sloth and the red howler monkey, to evaluate their patterns of habitat use and co-occurrence using a Bayesian occupancy modelling (Royle and Dorazio, 2008). Specifically, we (i) compare habitat use by jaguars in sites embedded in islands and the continuous forest, and (ii) evaluate whether the local relative abundance of both prey species influences the probability of habitat use by jaguars. We hypothesize that jaguars’ habitat use is similar in islands and continuous forest, and that prey abundances are strong predictors of it.

## Methods

### Study area

Our study region comprises the *várzea* forest, a floodplain forest ecosystem located at the confluence of the Solimões and Japurá rivers in Central Amazon (Fig. 1). The interfluvium at these rivers’ junction is protected by the Mamirauá Sustainable Development Reserve (IDSM, 2010). *Várzea* forests are seasonally flooded by nutrient-rich white-water rivers, whose average annual water level range is 12 m (Ramalho et al., 2009). The maximum water level is reached around June and its minimum between October and November (IDSM, 2010).

**Figure 1.**
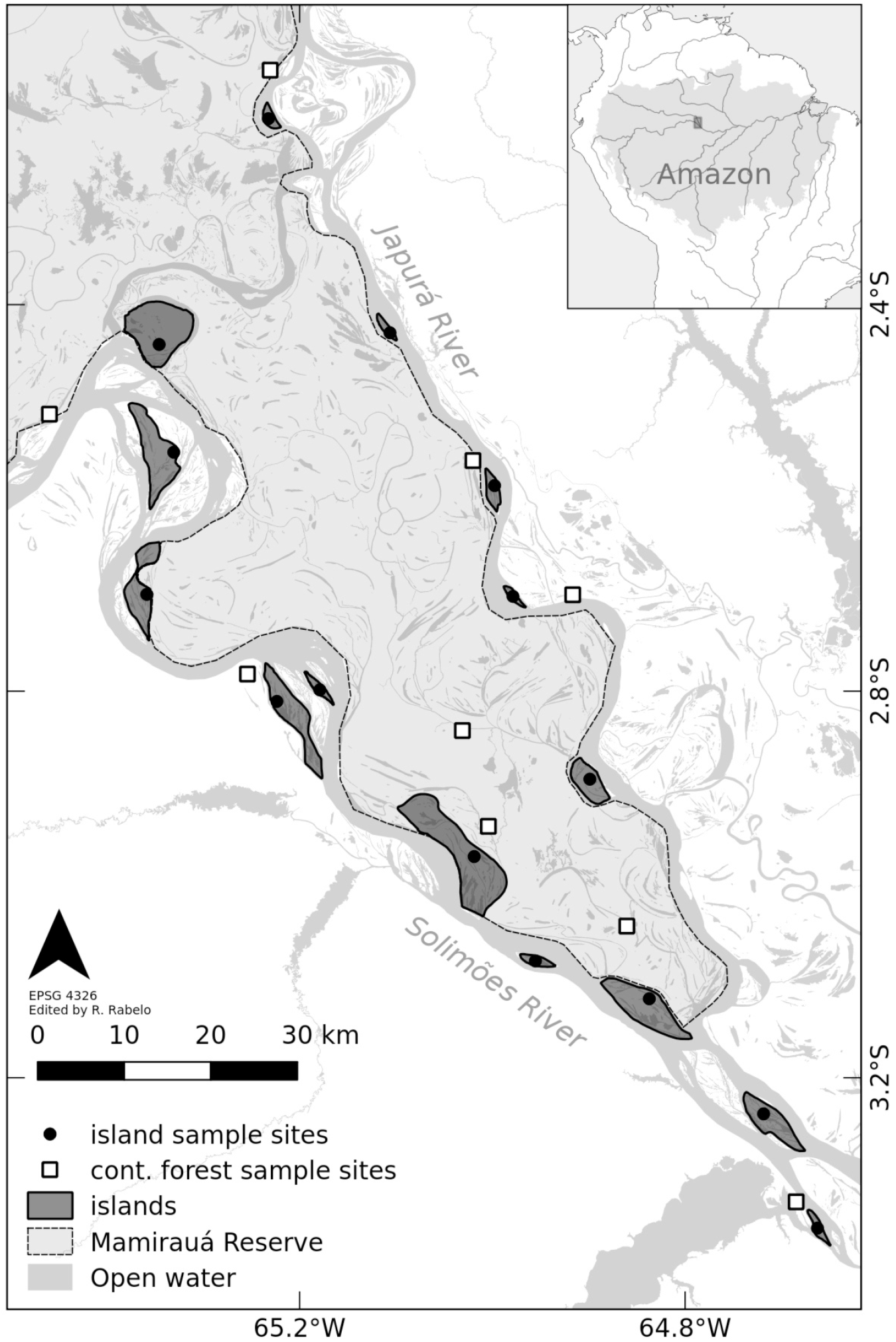
Map of the study region showing the distribution of sample sites in the Central Amazonia floodplain forest landscape.

River dynamics constantly modifies the spatial structure of these riverscapes by the erosion of margins and the transport and deposition of sediments (Peixoto et al., 2009). This process creates fluvial islands that emerge and can disappear in a few decades (Kalliola et al., 1991). Although some islands represent ephemeral habitat patches for mammals with long generation times, such as jaguars, these species often use them (Rabelo et al., 2019).

### Sampling design and data collection

We sampled 14 focal islands ranging from 151 to 3,625 ha and nine independent sample sites embedded in the adjacent continuous forest (Fig. 1). We chose islands (i) permanently surrounded by the water matrix (that is, even during the low-water season), (ii) whose edge was at least 2 km distant from the edge of the nearest sampled island to avoid sampling islands that are too close to each other, and (iii) ≥30 years-old (island age was determined using a historical series of Landsat Thematic Mapper satellite images) to avoid islands that are too ephemeral for our study species (jaguar generation time: ~7 years, de la Torre et al., 2018). Although we believe that jaguars can visit islands younger than 30-years old during their daily journeys, it is unlikely that these islands have an adequate forest structure to harbor arboreal mammals, as most islands younger than 30 years are dominated by pioneer vegetation and rarely present late-succession forest patches (Peixoto et al. 2009).

We sampled mammals along line-transects that are independent sample sites. Transect length on islands varied from 1.2 km to 11.6 km and it was directly correlated to island size (Pearson correlation: *r* = 0.94, *P* < 0.001), making island size an intrinsic characteristic of each sample. We established line-transects with the same length range within the continuous forest sample sites to survey species. Surveys consisted of quiet walks on trails by two trained observers at *ca*. 1.5 km/h following a standardized protocol (Peres, 1999). We carried out the surveys in the morning (0630–1130 h) and afternoon (1300–1700 h). We stopped surveying when it was raining. We recorded sloths and howler monkeys via sighting and vestiges (e.g., calls and feces) and jaguars via fresh feces and footprints (recorded signs were marked to avoid double detection). Although some researchers can argue that the other wild Amazonian big cat (puma, *Puma concolor*) could be responsible for the vestiges, no records of this species were obtained in an effort of 2,040 camera*day at Mamirauá Reserve (Alvarenga et al., 2018). Therefore, we assumed that there is no resident puma population in the reserve. We conducted four replicate surveys per transect (i.e., four occasions), separated by one to four days, during the low-water season (September to November) of either 2013 or 2014. We limited surveys to the low-water season to minimize potential seasonal effects on species detection. We were unable to visit all sample sites during a single low-water season due to logistical constraints.

### Data analysis

We assessed the pattern of jaguar occurrence across our sample sites using an occupancy modelling approach. This approach estimates the probability of a site being occupied/used (*ψ*) by a given species when its detection is imperfect; that is, when the detection probability is less than 1 (Mackenzie et al., 2006). Given that the non-detection of a species at a sample site results from either its true absence or the failure to detect it, repeated surveys (occasions) on multiple sample sites are used to estimate the detection probability (*p*) of a species conditional on occupancy. In our model, jaguar occurrence at a given site “*j*” of type “*k*” (i.e., island or continuous forest) is denoted as *O_jk_* (i.e., the true occupancy state: 1 if present, 0 otherwise), and is the outcome of a Bernoulli trial with probability of occupancy *ψ_jk_*,

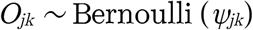

Similarly, the binomial detection/non-detection (1 = present; 0 = not detected) of jaguars (*D*) during a given occasion *“i”* and in a given sample site *“j”* are input in the form of an array *D_ij_*. Therefore, whether the species is detected during a given occasion in a given site is conditional on the occupancy state *O_jk_*, as follows:

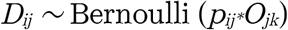

where *p_ij_* is the probability of jaguar detection during an occasion (survey) in a site.

We estimated both *ψ* and *p* parameters as linear responses to predictor variables using a logit link function as performed in a regular logistic model. Then, we added the line-transect length (values were standardized before running the model) into the model as a jaguar detection covariate; that is, we expected that the more we walked, the higher is the likelihood of detecting jaguar signs. We modeled the logit transformation of detection probability as follows:

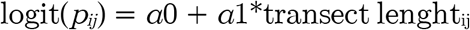

We also expected that transect length, which was positively correlated with island size, influences prey species counts. We estimated prey relative abundances (*λ_ij_* as linear responses to transect length using a log-link function as in a Poisson regression model as follows:

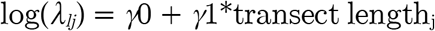

where *λ_jl_* is the relative abundance of prey species “*l*” (i.e., sloth or howler monkey) in site “*j*”. Finally, we considered that the occupancy probability *ψ_jk_* depends on the type of site “*k*” determined by *β*0 and on the relative abundances of sloths and howlers on site “*j*’ (counts of prey species were centered and scaled before running the model to compare their coefficients), in a logit transformation of a linear model as follows:

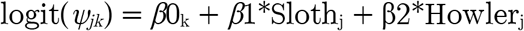

The full hierarchical model formulation is presented in Fig. S1. We implemented the model in a Bayesian framework using JAGS accessed via the software R, version 3.4.1 (R Core Team, 2017) using the package ‘rjags’ (Plummer, 2016) (see the R code available in Appendix S1). We used flat priors normally distributed with mean = 0 and variance = 100 for all model parameters, except for *SD* of *β*0, for which we used a gamma distribution with shape and rate equal to 0.1. We performed the estimation of posterior parameters with the Markov Chain Monte Carlo (MCMC) method using three parallel MCMC chains of 100,000 in length after discarding the first 10,000 steps of each as burn-in, and with a thinning rate of 100 steps. This combination of values ensured that all chains converged, i.e. essentially oscillated around the same mean parameter value (see Fig. S2) We report the posterior distribution of all estimated parameters as means and standard errors (*SE*), as well as the medians and the 2.5 and 97.5 percentiles, which are the Bayesian equivalent to the 95% confidence interval (highest posterior density in 95% – HPD95).

## Results

We obtained a total of 20 jaguar independent detections at 15 (63%) of the 24 sample sites during the four sampling occasions. Counts of both prey species were similar in islands and continuous forest (Fig. S3). Sloth counts per transect ranged from 0 to 8 individuals. They occurred at 13 (54%) sample sites. Sloth median counts per transect were 1 (first quartile (Q1) and third quartile (Q3) = 0 and 2.5, respectively) in islands and 0 (Q1 = 0; Q3 = 1) in continuous forest (Fig. S3a). Howler counts per transect ranged from 0 to 26 individuals. They inhabited 17 (71%) sample sites. The median value of howler counts per transect was 3 individuals (Q1 = 0; Q3 = 5.5) in islands and 5 individuals (Q1 = 2; Q3 = 6) in the continuous forest (Fig. S3b). The number of sites with predator-prey co-occurrences was 13 (54%) for sites shared by jaguars and howlers, and 11 (46%) for jaguars and sloths.

The mean jaguar detection probability across sites was *p* = 0.26 (HPD95: 0.15–0.39), and it was directly influenced by transect length (Table 1). Transect length was also a good predictor of the relative abundance of both prey species (Table 1; Fig. S4).

**Table 1.** Parameter estimates (link scale) from the hierarchical occupancy model for jaguar occurrence in Mamirauá’s floodplain forests in Central Amazonia. *γ* and *δ* are the coefficients (intercept and slope) of Poisson regressions for the effect of transect length on sloth and howler counts, respectively; *α* = coefficients (intercept and slope) of logistic model of jaguar detection probability *p*; *β* = coefficients (intercept and slopes) of logistic model of jaguar probability of occurrence *ψ*. See full model formulation in Fig. S1.

| Species | Parameter | Estimate (SE) | 2.5% | Median | 97.5% |
| --- | --- | --- | --- | --- | --- |
| Sloth | $\gamma_0$ | 0.21 (0.00) | -0.18 | 0.22 | 0.58 |
| | $\gamma_1$ | 0.36 (0.00) | 0.06 | 0.36 | 0.65 |
| Howler | $\delta_0$ | 1.33 (0.00) | 1.10 | 1.33 | 1.54 |
| | $\delta_1$ | 0.67 (0.00) | 0.52 | 0.67 | 0.81 |
| Jaguar | $\alpha_0$ | -1.05 (0.01) | -1.71 | -1.04 | -0.43 |
| | $\alpha_1$ | 0.33 (0.00) | -0.19 | 0.33 | 0.85 |
| | $\beta_0$[islands] | 3.27 (0.03) | 1.53 | 2.99 | 6.72 |
| | $\beta_0$[cont. forest] | 3.26 (0.03) | 1.53 | 2.97 | 6.70 |
| | $\beta_1$ (sloth) | 2.52 (0.03) | -0.66 | 2.37 | 6.44 |
| | $\beta_2$ (howler) | 2.52 (0.03) | -0.53 | 2.43 | 6.06 |

Jaguar probability of site occupancy was positively influenced by the abundance of both howlers and sloths (Table 1; Fig. 2a and b). Although both estimates were not significant at HPD95 (i.e., the HPD95 interval included the zero), we found strong evidence that sloth and howler abundances increase the probability of habitat occupancy by jaguars [likelihood estimates: p(*β*1 > 0) = 0.93 and p(*β*2 > 0) = 0.94, respectively]. We estimated a similar proportion of island and continuous forest sites used by jaguars (*ψ* = 0.75, HPD95: 0.36–1.00, Fig. 2c). Additionally, we found that the abundances of both sloths and howlers have similar effects on jaguar probability of occurrence (*β*1 - *β*2 = −0.002, HPD95: −4.38–4.56; Fig. 2d).

**Figure 2.**
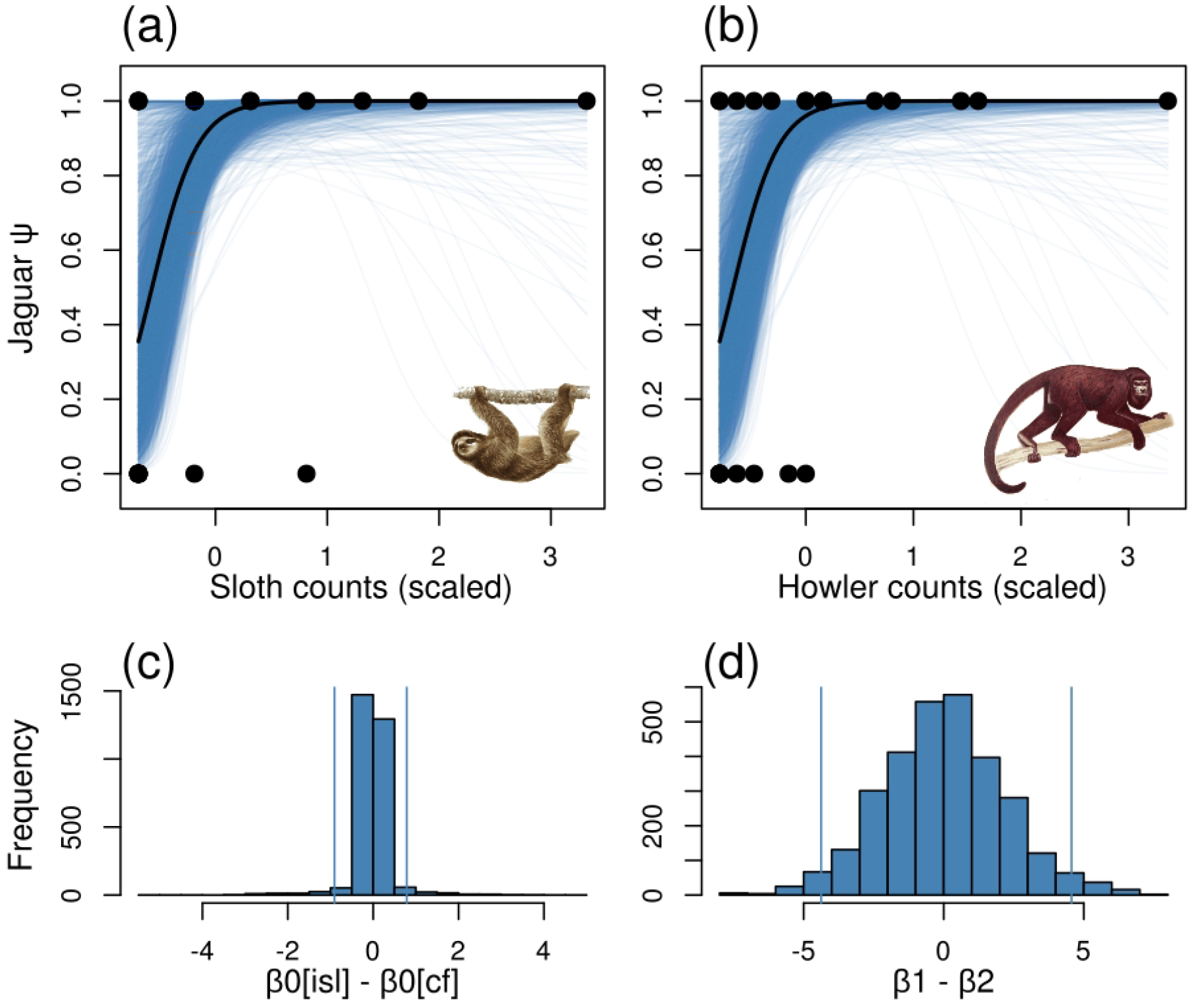
Predictors of jaguar occupancy across sample sites in Mamirauá’s floodplain forest in Central Amazonia. The logit transformation of jaguar probability of habitat occupancy (*ψ*) was modelled as a linear function of sample site type (i.e., embedded either in island or in continuous forest), and the abundance of prey species at the sample site. Jaguar mean probability of occurrence (black lines) increases markedly with increasing abundance of both sloths (a) and howlers (b). Blue lines represent all models fitted according to posteriors estimates and higher density of lines indicates the area with higher model confidence. There is strong evidence that jaguar probability of occurrence does not differ between sites embedded in islands or continuous forest, as shown by the distribution of differences between *β*0 for islands and for continuous forest (c). Both sloths (*β*1) and howlers (*β*2) counts showed similar effects on jaguar probability of habitat use (d). Illustrations from Stephen Nash (howler) and John Oriszo (sloth).

## Discussion

Here we provide the first estimates of jaguar occupancy patterns using sign surveys along line-transects. We recorded jaguar feces and footprints on trails of different lengths, accounting for imperfect detection and considering transect length as a detection covariate. We found an average probability of jaguar detection that is similar to previous estimates based on camera trap surveys, and a higher probability of habitat occupancy than most reported estimates (Table 2). Jaguar populations of Amazonian *várzea* forests have high densities and abundances (Ramalho, 2012), which allow to expect that they also have high detectability and habitat occupancy. However, we acknowledge that care needs to be taken in interpreting comparisons between survey estimates from studies differing in survey method (Table 2) and other factors (e.g., habitat type and quality, sampling effort, location, etc.). Despite that, we suggest that sign surveys along line-transects are useful for assessing jaguar occupancy patterns as they do not require expensive equipment. We urge for such comparative studies to assess the level of correlation between their estimates.

**Table 2.** Comparisons of jaguar mean probabilities of habitat occupancy (*ψ*) and detection (*p*) among studies.

| Survey method | $\psi$ | $p$ | Location | Reference |
| --- | --- | --- | --- | --- |
| Line-transects | 0.75 | 0.26 | Mamirauá Reserve, Northern Brazil | This study |
| Interviews | 0.57 | 0.28 | Atlantic Coast, Nicaragua | Zeller et al. (2011) |
| Camera traps | 0.77 | 0.12 | Madre de Dios Department, Peru | Tobler et al. (2015) |
|  | 0.69 | 0.18 | Iguaçu National Park, Southern Brazil | Silva et al. (2018) |
|  | 0.54 | 0.21 | Emas National Park, Central Brazil | Sollmann et al. (2012) |
|  | 0.42 | 0.26 | Magdalena River Valley, Colombia | Boron et al. (2018) |
|  | 0.42 | 0.03 | Iwokrama Forest, Guyana | Roopsind et al. (2017) |

We found no difference in jaguar probability of habitat use between island and continuous forest samples sites. Jaguars’ probability of occurrence tends to be higher in core areas with high forest cover and close to water (Boron et al., 2018; Silva et al., 2018; Sollmann et al., 2012; Zeller et al., 2011), and that experience low levels of anthropogenic disturbance (Roopsind et al. 2017; Silva et al. 2018). Although river dynamics constantly changes the spatial structure of Amazonian riverscapes (Peixoto et al., 2009), our study *várzea* landscape of Central Amazonia is characterized by high forest cover and low human density. Additionally, jaguars move far and easily among distinct environments (Morato et al., 2016) and spontaneously cross rivers and lakes (Holt, 1932). Therefore, we expect that the aquatic matrix that surrounds fluvial islands do not hamper jaguar dispersal across rivers to reach these well-preserved island forests.

We found that the relative local abundance of prey species were good predictors of jaguar habitat use patterns. Prey density was the best predictor of habitat use by tigers (*Panthera tigris*) in a metapopulation also surveyed via signs along trails in southern India (Karanth et al., 2011). Likewise, the local occurrence of prey species was more important than the distance to water or forest structure to explain jaguar habitat use at the Calakmul Biosphere Reserve in southeastern Mexico (Booker, 2016). Given that both prey species are equally abundant in islands and continuous forest sites (Fig. S3), we suggest that foraging for prey plays a critical role in jaguars’ decision to use fluvial islands.

We found that both prey species had similar effects on jaguar probability of habitat use, despite the lower range of sloth counts per site. Whereas jaguars inhabiting nonflooded habitats feed mostly on terrestrial prey (González and Miller, 2002), Mamirauá’s jaguars rely heavily on caimans and their eggs (Torralvo et al., 2017) and on arboreal mammals (e.g., sloths, howlers and lesser tamanduas [*Tamandua tetradactyla*]), as terrestrial prey is often missing in flooded forests (Ramalho, 2012). On the other hand, caimans (Da Silveira et al., 2008), sloths and howlers (Queiroz, 1995) occur at high densities in these forests, facilitating predator-prey encounters. Despite the aforementioned lower abundance of sloths in our flooded forests, their solitary life-style and lower mobility and defense ability, make them an easy prey, thereby potentially explaining the equal effect of both prey species on jaguar habitat use and the importance of sloths to its diet.

In sum, we show that the aquatic matrix surrounding fluvial islands does not hamper jaguars to use them. The importance of the abundance of both prey species as predictors of jaguar habitat use is compatible with the hypothesis that they often visit these islands as part of their prey search strategies. Additionally, we showed that sign surveys along trails is an alternative method for assessing the pattern of habitat use by jaguars in Amazonian *várzea* forests, although further and proper tests of its effectiveness must be conducted (e.g., Ancillotto et al., 2018). Finally, although we conducted this study during the low-water season, we believe that the importance of these arboreal mammals in the diet of jaguars is even higher during the high-water season, when jaguars spend most of their time in the forest canopy.

## Acknowledgements

The Instituto de Desenvolvimento Sustentável Mamirauá (IDSM-OS/MCTI) funded this research. RMR received fellowships from National Council for Scientific and Technological Development (CNPq, #131077/2014-7 and #142352/2017-9). We thank the field assistance of Jonei Brasil, André Neves and Jairo Neves. The Infrastructure and Logistics and Administration Teams of the Instituto de Desenvolvimento Sustentável Mamirauá supported our **fi**eld activities. We thank Cristian Dambros for providing valuable help on model formulation and implementation and Daniel Rocha for insightful contributions on the first draft. We also thank Luca Luiselli, Emiliano Mori and two anonymous reviewers for their critical evaluation of the manuscript.

## Supporting Information

**Figure S1.** Model formulation.

**Appendix S1.** R code for model implementation.

**Figure S2.** Chain convergence.

**Figure S3.** Prey species counts on island and continuous forest sample sites.

**Figure S4.** Effects of line-transect length on counts of prey species.

