## Supporting Information for "Prey abundance drives habitat occupancy by jaguars in Amazonian floodplain river islands" (from another version of this preprint: version 2, 2019-04-11)

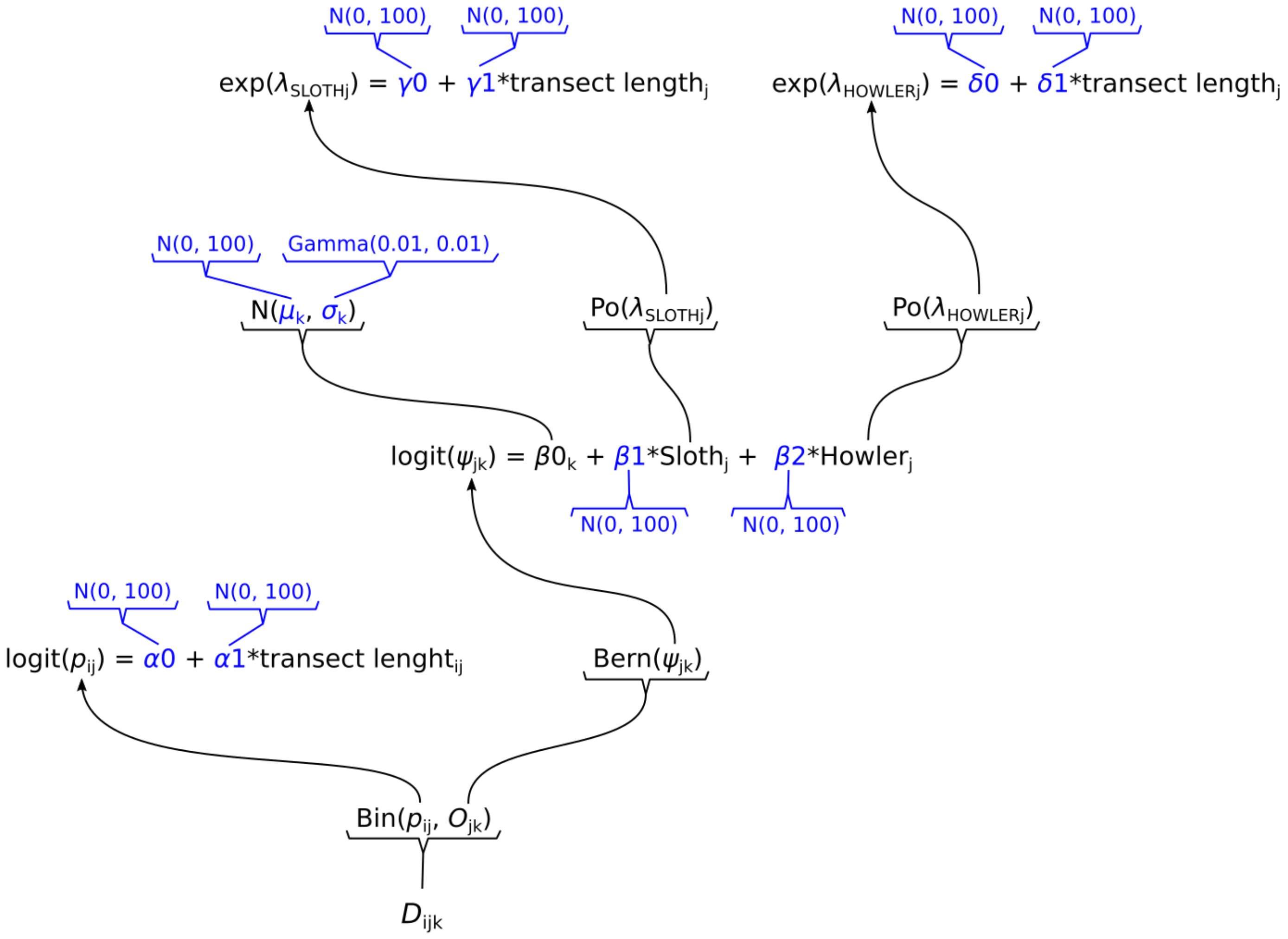


**Figure S1.** Representation of the Bayesian hierarchical occupancy model formulation used in the study. Blue coefficients represent the estimated model parameters, for which we provided flat priors coming from normal distribution with $\mu$ = 0 and $\sigma$ = 100, except for the $\sigma$ of $\beta 0$, in which we used a gamma distribution with shape and rate equals to 0.1.

**Appendix S1.** R code for model implementation

```
###############
## Habitat occupancy by jaguars in Amazonian fluvial islands
## R. Rabelo et al.
##
## Script created by Rafael Rabelo (last edited in 2018/Sep/04)
###############

library(rjags)

###############
## Model formulation

model<- "
model{
## Occupancy and detection modelling
for(i in 1:nsites){
  for(j in 1:noccasions){
    onca[i,j]~dbinom(det[i] * occ[i], 1)
  }

  ## Detection as a function of transect length
  detLinear[i]<- e0 + e1*TRANS[i,1]
  det[i]<- exp(detLinear[i])/(1 + exp(detLinear[i]))

  ## Occupancy as a function of site type (island/cont. forest),
  ## and of sloths and howlers abundance
  psiLinear[i]<- b0[t[i]] + b1*SLO[i] + b2*HOW[i]
  psi[i]<- exp(psiLinear[i])/(1 + exp(psiLinear[i]))

  occ[i]~dbinom(psi[i], 1)

  ## Abundance of preys as a function of transect length
  SLO[i]~dpois(lbda.SLO[i])
  lbda.SLO[i]<- exp(c0 + c1*TRANS[i,1])

  HOW[i]~dpois(lbda.HOW[i])
  lbda.HOW[i]<- exp(d0 + d1*TRANS[i,1])
}

  ## Loop for site type
  for(type in 1:2){
    b0[type]~dnorm(mut, 1/sdt)
  }

## Priors
e0~dnorm(0,1/10)
e1~dnorm(0,1/10)
```

```
mut~dnorm(0,1/10)
sdt~dgamma(0.1,0.1)
b1~dnorm(0,1/10)
b2~dnorm(0,1/10)
c0~dnorm(0,1/10)
c1~dnorm(0,1/10)
d0~dnorm(0,1/10)
d1~dnorm(0,1/10)
}
"

## Exporting model
writeLines(model, con="modelPredPrey.jags")


###############
## Model implementation

## Creating the data. Here I need to provide the
## names of each data as they appear within the
## model code, i.e., the diagram.
data<- list(onca=onca,
            nsites=nrow(onca),
            noccasions=ncol(onca),
            t=type,
            TRANS=trans,
            SLO=sloth,
            HOW=howler)

## Burn-in
ppModel<- jags.model("modelPredPrey.jags",
                      data=data,
                      n.chains=3,
                      n.adapt=10000,
                      inits = list(occ=rep(1,24)))

## Parameters estimates after chains convergence
ppResu<- coda.samples(ppModel,
                      variable.names=c("b0","b1","b2",
                                       "e0","e1","c0","c1",
                                       "d0","d1"),
                      n.iter=100000,
                      thin=100)

## transforming the results into a matrix
resu.mat<- as.matrix(ppResu); head(resu.mat)
## END
###############
```

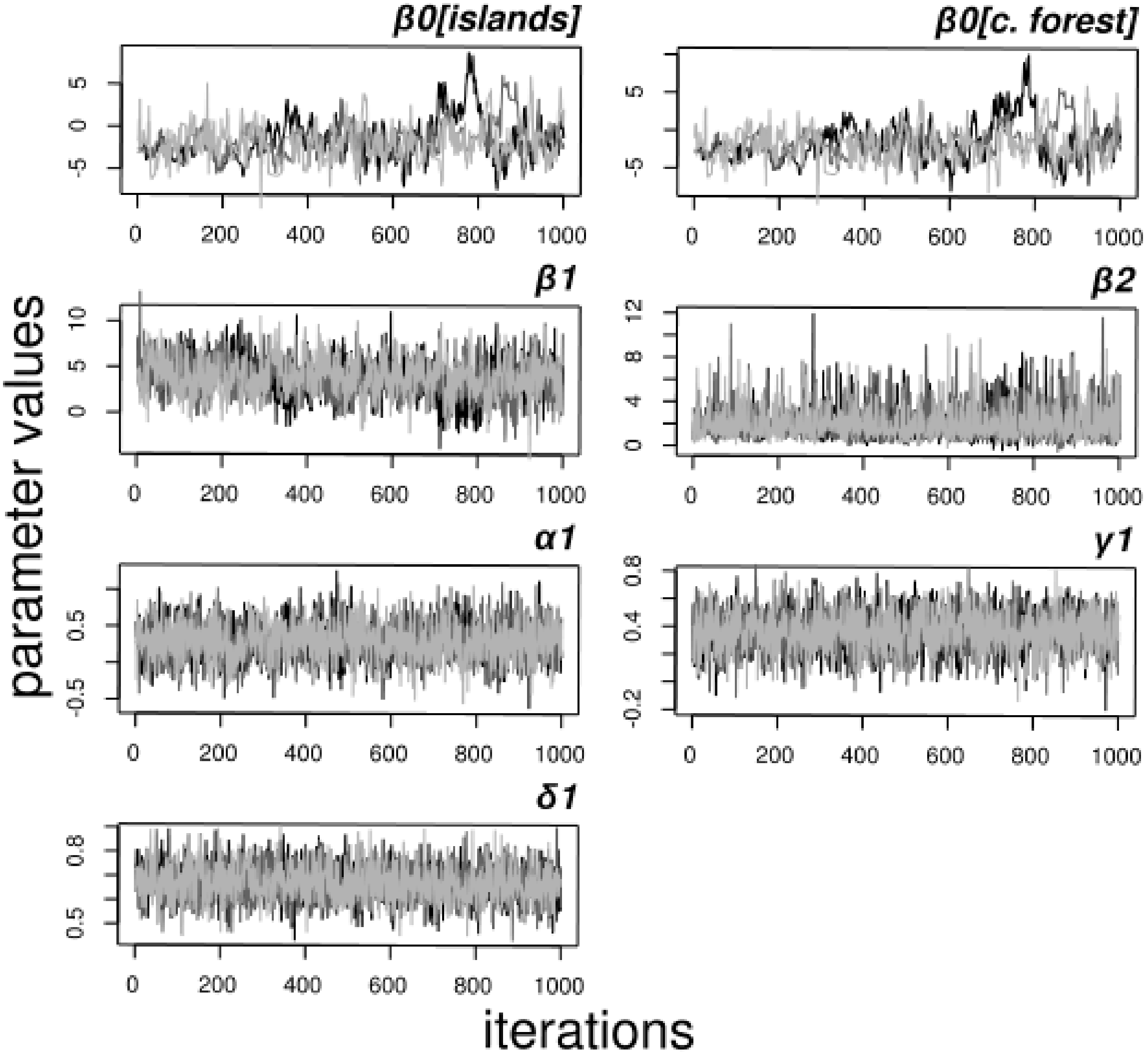


**Figure S2.** Chains convergence. Variation of parameter estimates along the iterations of Markov chains, after removal of burn-in steps. Light gray, dark gray and black lines represent each of the three parallel chains. It is possible to observe the chain convergence in parameter estimates along iterations for all parameters.

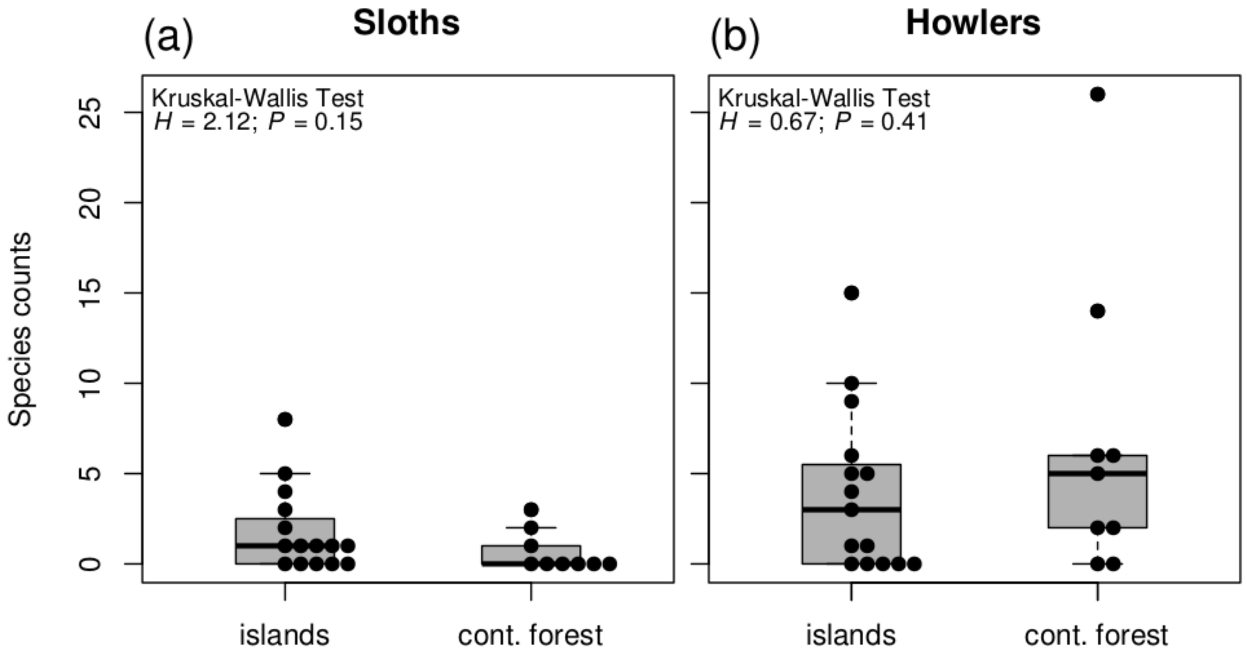


**Figure S3.** Prey species counts on islands and continuous forest sample sites. Each point represent a sample site (line-transect), which is embedded either on island or continuous forest (*x*-axis), and record the number of counted individuals (*y*-axis) on that sample site. Grey box-plots show the distribution of observations. According to Kruskal-Wallis non-parametric test, the number of counted individuals on line-transects did not differ significantly between islands and continuous forest sample sites for both sloths (a) and howlers (b).

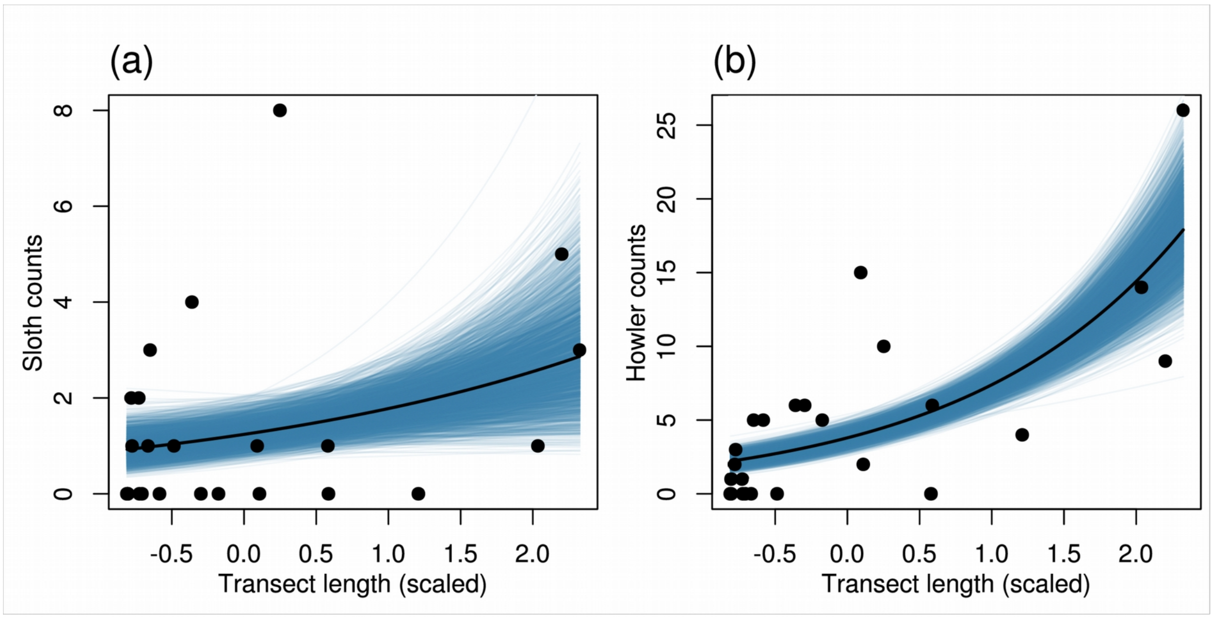


**Figure S4.** Effects of line-transect length on counts of prey species in Mamirauá's floodplain forest in Central Amazonia. Prey counts were used to represent the relative local abundances of each species. The longer the walk, the higher the number of records of both sloths (a) and howlers (b). The black line represents the median exponential response of prey abundance as predicted by a GLM with Poisson distribution formulated in a Bayesian framework. Blue lines represent all models fitted according to posteriors estimates.
